# Identity and compatibility of reference genome resources

**DOI:** 10.1101/2021.03.15.435425

**Authors:** Michał Stolarczyk, Bingjie Xue, Nathan C. Sheffield

## Abstract

Genome analysis relies on reference data like sequences, feature annotations, and aligner indexes. These data can be found in many versions from many sources, making it challenging to identify and assess compatibility among them. For example, how can you determine which indexes are derived from identical raw sequence files, or which annotations share a compatible coordinate system? Here, we describe a novel approach to establish identity and compatibility of reference genome resources. We approach this with three advances: First, we derive unique identifiers for each resource; second, we record parent-child relationships among resources; and third, we describe recursive identifiers that determine identity as well as compatibility of coordinate systems and sequence names. These advances facilitate portability, reproducibility, and re-use of genome reference data.

**Availability:** https://refgenie.databio.org

## Introduction

Reference genome assemblies are representations of a genome (1–5) that are the basis of many prerequisites of genome analysis, such as alignment indexes (6–9) and feature annotations (10–12). Several tools under development aid in organizing and sharing such genome-related data (13–16), including our recent software called *refgenie* (17). Refgenie is a genome resource asset manager that provides two ways to obtain genome assets: Users may *pull* pre-built assets from a remote server, or *build* equivalent assets locally. This flexibility increases interoperability of tools that rely on genome assets; however, it also raises challenges with identity and compatibility of these assets.

One common challenge is identifier mismatches. Relying on simple human-readable identifiers such as “hg38” means two users may refer to different things with the same identifier. As a case in point, there are many variations of the human genome that are all referred to in different analysis as “hg38” or “GRCh38”. This leads to compatibility issues that incur the wrath of bioinformaticians everywhere. A step toward solving this problem is to use unique identifiers that unambiguously identify a particular assembly, such as those provided by the NCBI Assembly database (4); however, this approach relies on a central authority, and therefore can not apply to custom genomes or assets.

Another weakness of centralized unique identifiers is that they are insufficient to *confirm* identity, which must also consider the content of the genome. For example, if someone makes a minor adjustment to reference data content, but continues referring to it with the centralized identifier, this can lead to reproducibility issues. To ensure that assets from different locations are identical not only in name, but in content, we require a more substantial way to uniquely identify *and confirm identity of* both assets and genomes. The situation is further complicated by assets that are derived from other assets. For example, a *bowtie2 index* is derived from a *fasta file*; trading around bowtie2 indexes without the underlying fasta asset can lead to downstream analysis incompatibilities. To solve this problem requires a way to record not just the identity of genomes, but the relationships among assets that are derived from them.

A method that is capable of confirming both the identity of and the relationships among assets solves these challenges; but what if we only need to confirm that a coordinate system is compatible? This is a less stringent comparison because it does not require identical genomes, but a more nuanced comparison among them. A common example is sharing feature annotation data across related genomes that do not necessarily have identical sequences, but do have an identical coordinate system. Establishing compatibility in this sense requires a more detailed comparison between the assets which cannot be accomplished with only unique identifiers and relationships. This requires capacity to assess not just *identity*, but *compatibility* between non-identical assets.

Recent advances partially address some of these challenges. First, refget (16) computes identifiers for a sequence from the sequence itself, and provides a lookup database to retrieve the sequence given its identifier. Refget thus provides a globally unique, content-derived identifier and retrieval system for raw sequences. A similar approach is also taken by the Variant Representation Specification (VRS) for identifying genetic variants (18). The tximeta package (19) similarly identifies transcriptomes based on similar sequence identifiers. But no existing approach provides a way to establish identity, relationships, and compatibility among genomes and arbitrary assets derived from them.

Here, we address each of these issues. Our approach can guarantee identity, relationships, and compatibility among reference genome assets, which we have implemented in our refgenie software. Refgenie accomplishes this with 3 concepts: First, for each asset, it computes unique asset identifiers that are derived from the assets themselves; Second, it records which parent assets were used to create each derived asset. Third, it employs a genome identifier system that allows it to not only establish the identity of a genome, but also to quickly compute multiple levels of compatibility between them. Together, these tools improve the interoperability and reproducibility of analytical pipelines that rely on reference genome assembly assets.

## Results

### Identity: Unique asset-derived identifiers

Refgenie asset keys are human-readable, which is great for humans, but can lead to name collisions; for instance, how can a user be sure that the *bowtie2 index* keyed at one location is the same as another? In a closed system where all assets are downloaded from a single server, this is not a problem; however, the refgenie system is flexible, allowing multiple servers, building custom assets locally, and human-readable identifiers that give the user total control. This makes refgenie flexible and powerful, but also means that identity cannot be guaranteed by name alone (Fig.1A).

**Fig. 1:**
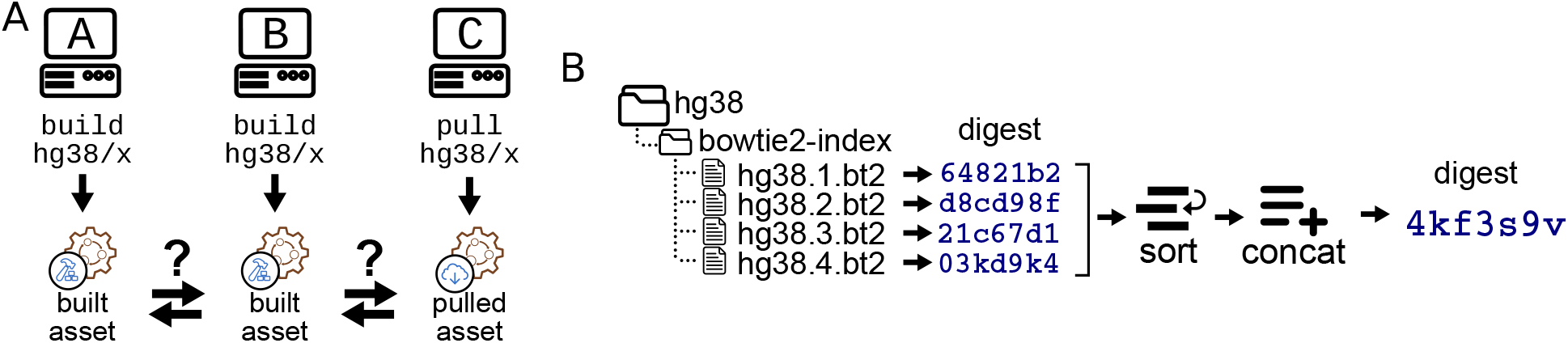
Unique asset-derived identifiers. A) Assets built on different systems using the same human-readable identifier may not be identical. Refgenie requires a way to establish the identity of assets on different servers. B) Refgenie’s unique asset-derived identifiers, such as shown here for a bowtie2 index, work by calculating a digest of each individual file in the asset, sorting these digests, concatenating them, and then calculating a final digest, which is the unique identifier for the asset. The files in this example are the 4 bowtie2 Burrows-Wheeler index files for hg38, which is an arbitrary example asset. Digests are shortened for illustration.

To address this issue, Refgenie requires a unique identifier for each asset. Critically, these identifiers must be computable for arbitrary, custom assets rather than created by a central authority, so they must be derived *from the assets themselves*. Furthermore, refgenie makes no assumptions about the data types of assets, so the identifiers must be compatible with any kind of data.

Refgenie accomplishes this with a simple hashing algorithm: we take all files in the asset folder, calculate the md5 digest on each file independently, lexographically sort the digests, and then calculate the md5 digest on the resulting list (Fig.1B). This is a straightforward method to derive a digest for a set of arbitrary files and is thus compatible with any type of asset. This identifier is automatically computed by the `refgenie build` process, thereby assigning a unique identifier to every asset.

The refgenie command-line interface (CLI) allows users to retrieve this identifier using the `id` command. For example, `refgenie id hg38/bowtie2 index:tag` would return the unique identifier for the specified asset. These identifiers establish an automated way to identify any possible asset, and can also be re-computed to confirm the true identity of an asset, regardless of its human-readable identifier. Refgenie relies on these globally unique and reproducible identifiers to refer to assets uniquely. This approach establishes universal identifiers that allow users to confirm asset identity across systems.

### Consistency: Recording asset relationships

One of refgenie’s strengths is *derived assets*: that is, assets that are easily built from other assets. For example, if a user has a `hg38/fasta` asset, then building a `hg38/bowtie2 index` asset requires no further inputs, as refgenie will automatically use the existing fasta asset to build the index. This is convenient for the user, but it can also lead to a potential conflict if a user then tries to `pull` an asset for the same genome that was derived from a different parent. For example, perhaps the user issues `refgenie pull hg38/bwa index`. The critical question is this: was the fasta file that was used to create the bwa index on the server identical to the one the user has tagged as ‘hg38’ locally? If not, the pull should fail. The only way to guarantee that derived assets have identical parents is to record the relationships among them (Fig. 2A).

**Fig. 2:**
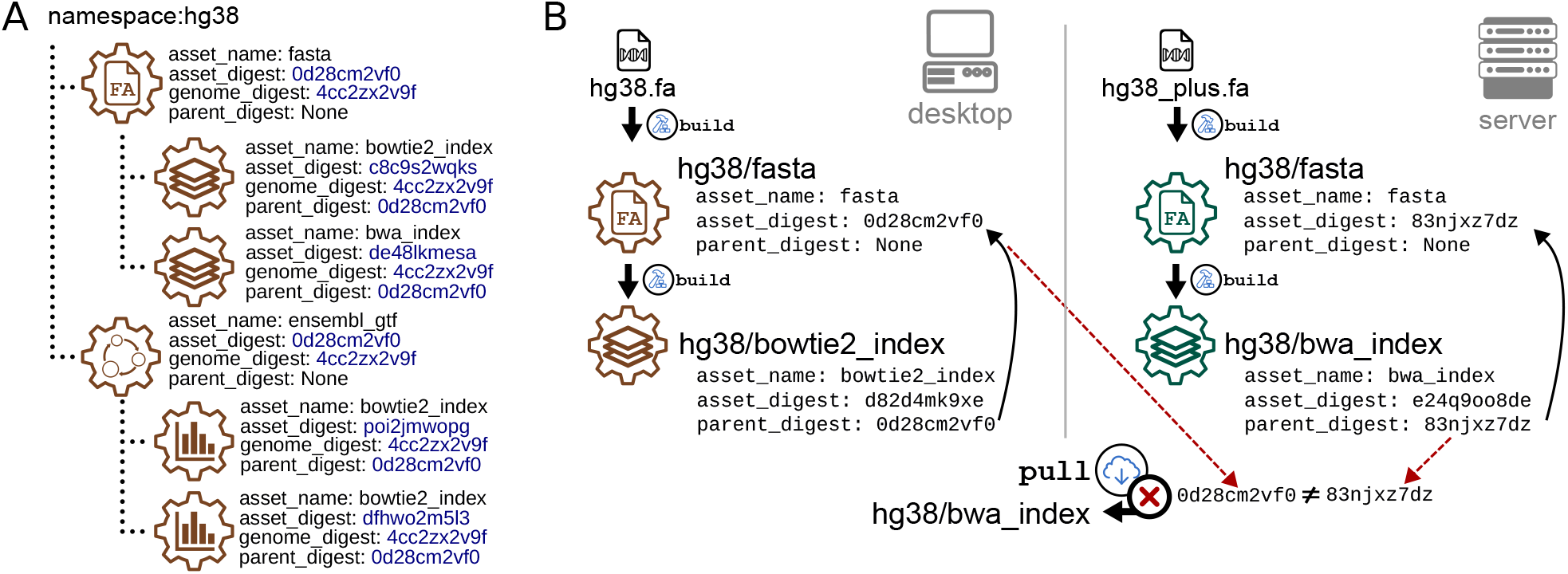
Recording relationships ensures compatibility of derived assets. A) Derived assets naturally form a parent-child structure. Refgenie records parent-child relationships by storing the unique identifier of all parent assets. B) For assets that can be built from other assets, we require a way to ensure that all parent assets match when using either build or pull to obtain the asset. Derived assets must be derived from the same parent assets to be compatible. For assets with multiple parents, all identifiers must match.

To solve this problem, refgenie `build` records not only the unique asset identifier, but also parent-child relationships. For example, the remote refgenie server entry for the `hg38/bowtie2 index` asset retains a pointer to the `hg38/fasta` asset that was used to build it (Fig. 2B). You can think of this as *each built asset remembers the unique identifier (not the human-readable identifier) of any assets used to build it*. Most assets have only a single parent, but refgenie allows assets to have multiple parents. When the `pull` command is issued, the CLI checks the parent identifiers against the local ones. If any digests do not match, refgenie aborts the pull request, preventing a user from mixing assets that have been derived from different parents. Without this check, the user could introduce an inconsistency because pulled assets were not built from the same input as built assets.

To make this procedure complete, refgenie stores the parent-child relationships and the CLI makes sure that these relationships are kept intact when users remove, re-tag, or rename assets. The digest check is fast because it does not require pulling the parent asset in its entirety; only checking for its unique identifier. To enable this, the refgenie server presents this information as an API endpoint. If a user pulls a derived asset when the parent does not exist locally, refgenie will populate the parent asset digest in the config file. In a sense, the first stage of pulling a derived asset from an unknown parent asset “locks” the parent asset, preventing pulling further assets from other sources that claim to be derived from the same parent, but are not.

### Compatibility: decomposable genome identifiers

Computing unique asset-derived identifiers plus storing asset relationships together allow us to record and compare asset identity and assure a consistent lineage of derived assets. This solves many challenges that require strictly identical assets. For example, to reproduce the result of a bowtie2 alignment requires an identical bowtie2 index asset. An index built from a fasta file that is identical in sequence content, but differs in identifier names or order will yield a different result. The strict asset-derived identifiers can ensure this level of reproducibility. Storing the relationships extends this assurance to derived assets, making it possible to ensure that they share identical parents.

However, many analyses require less stringent comparison: simple compatibility between assets that are not necessarily identical. For instance, a motif enrichment analysis is strictly tied to a specific sequence, but the order of the original sequences may not be relevant for the results to be comparable. As a result, using a fasta asset with identical sequence but different order would not be a problem. Some analysis require even less strict requirements. For example, say analysis *annotate regions* reads an aligned bam file, and annotates it using feature annotations on the *hg38* coordinate system. This analysis requires the reads to share the coordinate system of the annotation, but it does not require a specific reference assembly *sequence* at all. In this case, we would like to confirm that the given *bowtie2 index* as-set *is compatible with* the feature annotation coordinate system. To fulfill this requires only that it shares a coordinate system, not that the exact sequence matches. In short, sometimes we do not require strict identity, but a more detailed comparison that may ignore order, sequence names, or other attributes. We therefore seek to distinguish between the comparison of *is compatible with* vs *is identical to*.

A system that relies only on unique identifiers cannot make this fine-grained of a comparison because it requires directly comparing not just the identifiers, but the *contents* of the assets of interest. Since this task must consider the contents of an asset, it is impossible to come up with a universal solution that works on any data type the way our generic asset-derived identifiers do. To establish compatibility related to genomic features such as nucleotide sequences, sequence names, genome membership, order, and length, we must therefore develop a more specific solution for this particular use case. To solve this problem, refgenie relies on a novel concept we refer to as *decomposable identifiers*.

#### Sequence collection identifiers

Refgenie’s approach is based on the refget protocol for identification and retrieval of sequences (16). In refget, DNA sequences are hashed to create a unique identifier that is stored in a database and can be used to retrieve the original sequence (Fig. 3A). Refget identifiers are specialized to DNA or protein sequences, and it adds a critical component of allowing *lookup* of the underlying data given the identifier. Lookup is not necessary for the identity and provenance objectives described above; however, it becomes important for the *compatibility* question, which requires asset content to ask a more fine-grained comparison question. Nevertheless, the current refget protocol only partially fulfills refgenie’s need because refget only accommodates individual sequences, and also does not allow for compatibility comparisons. To answer the compatibility questions, we devised a new digest procedure that extends the refget protocol in two ways: First, we extend to annotated sets of sequences; and second, we add the *length* of the sequence as a metadata component to the string to digest. We refer to this as a decomposable identifier because after a retrieval, the original object is not a simple sequence, but a tuple that can then be decomposed into constituent parts.

**Fig. 3:**
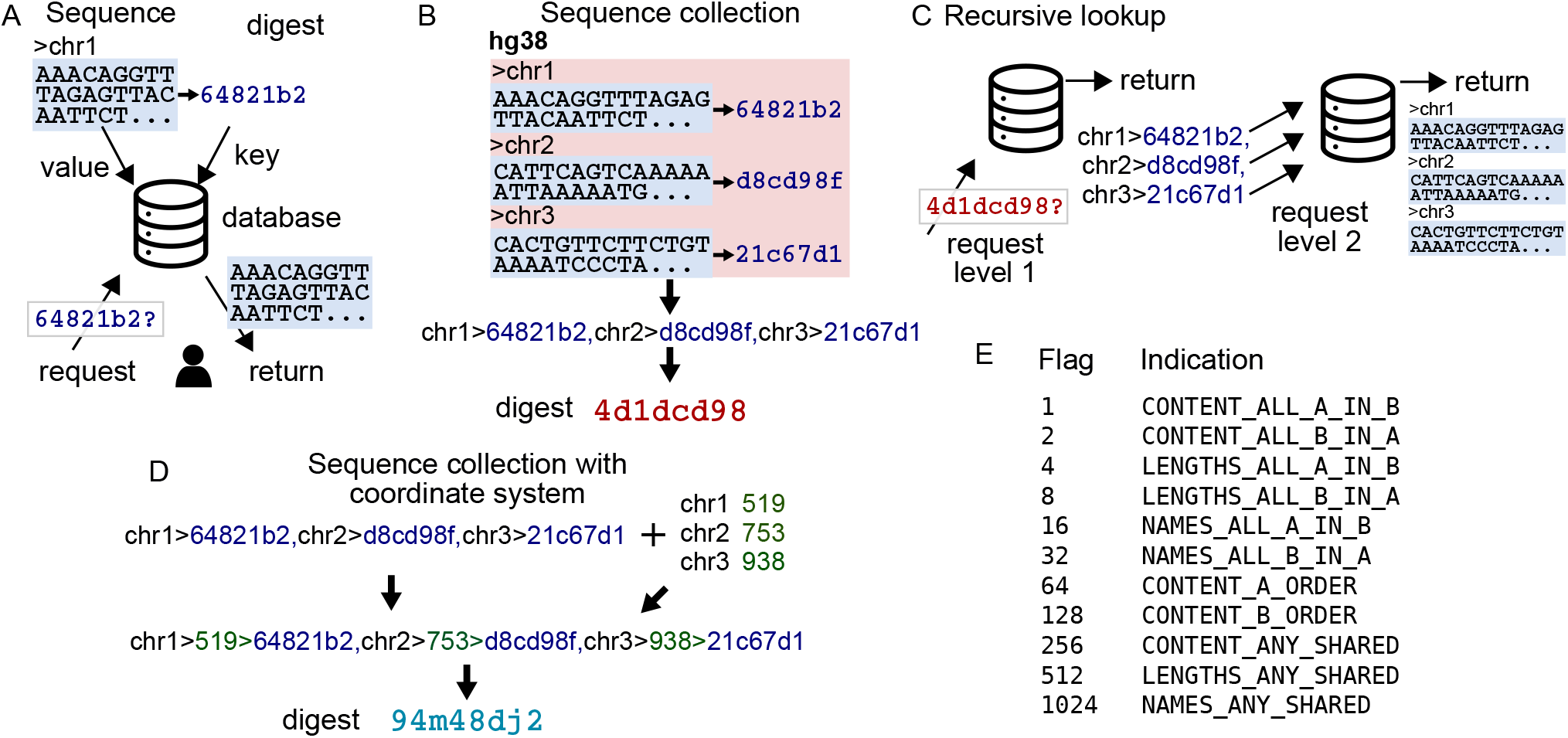
Decomposible recursive unique identifiers. A) The refget protocol uniquely identifies and retrieves DNA sequences. First, a sequence is hashed to yield a digest, which is used as a unique identifier to store the sequence in a database. A request using the unique identifier returns the original sequences. Refget digests uniquely identify a DNA sequence and provide a way to retrieve the sequence using the unique identifier. B) A sequence collection digest is made by first computing refget digests for each sequence, concatenating them, and computing a digest on the result. C) Sequence collection digests can be used to retrieve the sequence collection recursively. In the first step, the string of digests is returned; each sequence digest can then be used to retrieve its sequence, finally yielding the sequence collection. D) By adding the sequence lengths to the digest string, a new string and digest can be made that allows retrieving sequence names, lengths, and digests. E) A table of flags provide a way to quickly indicate the relationship between two sequence collections.

##### 1. Annotated collections of sequences

We first hash the sequences themselves, then we concatenate them with delimited sequence identifiers, and compute the digest of the resulting string (Fig. 3B). The dual-delimited string uses one character, here “,”, to delimit items (sequences), and another character, here “>“, to delimit the attributes of the items (names and sequence digests).

Since one of the attributes of the sequences is itself a digest, the final unique digest is a digest of digests. This recursive approach accomplishes several goals. First, it satisfies the goal of creating a checksum that can be used to confirm identity of collections of sequences (for example, fasta files). Second, it also allows the ability to do more detailed comparison between two collections; for example, we can check the sequence-level checksums to see if two fasta files have the same sequences, but in different order or with different names.

In this toy example, a lookup of digest `4d1dcd98` would return this string:

~~~
chr1>64821b2,chr2>d8cd98f,chr3>21c67d1
~~~

This string allows us to compare the names and content of this against another sequence collection. For example, say another lookup returned this string:

~~~
chr1>64821b2,chr3>21c67d1,chr2>d8cd98f
~~~

In this case, a quick comparison would identify that these collections have identical sequences and names, but in different order. In similar way, a quick comparison could identify if sequence *content* matches but *names* do not, or if one collection is a subset of another, or if two collections are completely different. These comparisons are all very efficient because no actual sequences are compared, only names and digests for each sequence.

The recursion of lookups provides additional power for comparison (Fig. 3C). A single call to the function accepts a recursion parameter that allows refgenie to return the complete fasta file. This allows users to reconstruct a complete reference assembly given nothing but its unique digest.

##### 2. Adding sequence lengths for coordinate system identifiers

The decomposable sequence identifier concept has solved most of the problems we outlined earlier; we can now compare sequences for differences in order, sequence identifier, membership, etc. However, there is still one common scenario that this does not accommodate: compatibility of coordinate systems. A coordinate system can be defined as a set of named sequences with lengths. For example:

~~~
chr1 519
chr2 753
chr3 938
~~~

A set of sequences by definition must have a coordinate system, but a coordinate system does not specify sequences.

We seek a system that will allow us to confirm that two assets use the same coordinate system, even if they use a completely different set of sequences. This question cannot be easily answered using the sequence collection digests alone; it requires retrieving the original sequences to compute their lengths. Because this compatibility question is a critical and frequent query, instead of requiring this additional computation, refgenie adds the lengths into the collection digest string source (Fig. 3D). To do this, we simply prepend the sequence length to the item string before the final digest. Now, when a sequence collection digest is used for lookup, we return 3 attributes of each sequence instead of 2: the *name, length*, and *sequence digest*. This addition allows us to rapidly make compatibility comparisons at the coordinate system level. Given two strings in this format, without needing to process the sequences themselves, we can quickly determine if two assemblies share a coordinate system. Refgenie needs to simply compare the names, lengths, and sequence digests; if the names and lengths match, then it is a reasonable to assume the coordinate systems are compatible. To enable this, refgenie stores the chromosome names, lengths, and sequence digests locally for any genome when a *fasta* asset is built or pulled. To make it simple to calculate the compatibility between two sequence collections, we have implemented a compare function, described next.

#### A component compatibility function

The compare function returns a flag, given two digests, *flag* = *f* (*digestA, digestB*). The flag returned is a binary indicator with bits set according to the relationship computed between the two given digests (Fig. 3E). This flag allows a user to easily test any of the possible compatibilities between the two digests using a simple logical operator. For example, to confirm that two sequence collections have identical sequence content, we use: *flag*&&1. To test if they have identical lengths, we use *flag*&&2. This flexible system allows the user to quickly identify compatibility across the whole spectrum, from use cases that require strict identity of identically named sequences in identical order, to flexible systems that require only a set of sequences that share sequence lengths.

Users can invoke the compare function directly from the command line using the `compare` command:

~~~
refgenie compare genomeA genomeB
~~~

## Discussion

Reference genomes, indexes, annotations, and other genome assets are integral to sequencing analysis projects. Refgenie provides a full-service management system that includes a convenient method for downloading, building, sharing, and using genome-based resources. As data availability increases, more tools are needed to provide for identity and compatibility of analysis. These tools are important piece in improving reproducibility of genomic analysis. With the updates described here, the refgenie system has been improved to provide a new way to establish genome asset identity, relationships, and compatibility. These improvements will make it easier to ensure reproducibility and track provenance of downstream analysis that is based on refgenie assets.

Being able to identify genomes is a critical task in bioinformatics. Here, we introduced a novel approach using recursive identifiers, which enable a new type of compatibility test that can establish multiple levels of compatibility among genome-related assets. This improvement will make it possible for downstream tools to more easily check compatibility of reference genome resources, improving their portability and reusability.

Development of refgenie is continuing, with several new features planned. Refgenie already handles any user-provided genome assemblies, such as a custom spike-in genome or a combined multi-species assembly. But an area for improvement will be the ability to specify custom assets. Currently, refgenie can only build a restricted set of assets, but we have started work on a more flexible approach with custom recipes so that users can add new asset types. A second area of rapid development is the potential to use refgenie to reference cloud resources. Currently, refgenie is built around retrieving remote assets for local use, but a future update could make it possible for a local refgenie client to provide cloud paths to unarchived assets, which could simply using refgenie in a pure cloud environment. Refgenie has already been adapted for easy use in Galaxy (20) and Snakemake (21) workflow systems, and we are interested in continuing to develop integrations with similar systems to make it easier for users to develop workflows that make use of refgenie reference data.

## License & availability

Refgenie consists of a series of Python packages that are all BSD2-licensed. Source code, documentation, and a list of active server instances can be found at refge- nie.databio.org.

## Author contributions

MS implemented the method and edited the paper. BX contributed to the implementation. NCS conceived of the study, contributed to implementation, and wrote the paper. All authors approved the paper.

## Acknowledgments

We thank the Global Alliance for Genomic Health (GA4GH) Refget working group for helping discussions on related topics.

